# DNA-functionalized Gold Nanorods for Targeted Triple-modal Optical Imaging and Photothermal Therapy of Triple-negative Breast Cancer

**DOI:** 10.1101/2022.02.12.480219

**Authors:** Suchetan Pal, Jaya Krishna Koneru, Chrysafis Andreou, Tatini Rakshit, Vinagolu K. Rajasekhar, Marek Wlodarczyk, John H. Healey, Moritz F. Kircher, Jagannath Mondal

## Abstract

Targeted imaging and therapy for triple-negative breast cancer (TNBC) in the perioperative period are believed to be imperative for better disease management and improved life expectancy. Still, they are not yet available in clinical settings, and only a few nanoparticle-based theranostic agents potentially offer these capabilities. Herein, we develop an innovative class of biocompatible triple-modality nanoprobes (TMNPs) that offer optical imaging using optoacoustic, fluorescence, and surface-enhanced Raman scattering (SERS), as well as photothermal therapy (PTT) with near-infrared (NIR) light. The TMNPs are fabricated by immobilizing positively charged NIR fluorophores on negatively charged DNA-coated gold nanorods (AuNR), then silica encapsulation. The DNA-based design allows the screening of commercially available positively charged NIR fluorophores for the optimum fluorescence and SERS signals. After the design optimization, we functionalize TMNPs with folate groups to target folate receptor1 (FOLR1)-overexpressing TNBC in vitro and in vivo. Our results reveal that TMNPs preferentially accumulate in the FOLR1 positive tumors in TNBC patient-derived xenograft mouse models and show excellent imaging capabilities with all three imaging modalities. Selective exposure of the tumor with NIR laser further shows efficient thermal tissue ablation without causing systemic toxicity. Collectively, TMNP holds great promise for real-time multiplexed imaging of cancer biomarkers and therapeutic capability.

## Introduction

Breast cancer, the most frequent malignancy in women, accounted for 0.68 million deaths worldwide in 2020.^1^ Specifically, the triple-negative breast cancer (TNBC) subtype is associated with poor prognosis in the patients, as it does not respond to hormonal and targeted chemotherapies.^2^ Surgical resection coupled with chemotherapy remains the most effective treatment regimen for nonmetastatic TNBC patients. Complete resection of the malignant tissue significantly reduces the chances of disease recurrence and improves life expectancy in such patients.^3^ However, complete resection of the entire tumor volume is challenging due to lack of visual contrast between cancer and healthy tissue. In addition, aggressive resection of excess healthy tissue surrounding the cancer tissue is often not reasonable, as critical blood vessels and nerves may be present in the tumor vicinity.^4^ To aid the surgeon in locating the tumor tissue, molecular imaging is performed, before or during surgery.^5^ Exogenous contrast agents are administered and tracked in a patient’s body via several imaging modalities, e. g., computed tomography (CT), magnetic resonance imaging (MRI), positron emission tomography (PET), and optical modalities (e.g., fluorescence, Cerenkov, optoacoustic, Raman scattering).^6-7^ Near-infrared (NIR) optical imaging modalities offer distinct advantages over non-optical modalities (e.g., PET, CT, MRI), such as less detrimental exposure to ionizing radiation, better sensitivity and resolution, and intraoperative imaging of several millimeters deep tissue.^8^ Nonetheless, small-molecule contrast agents infiltrate both cancer and healthy tissue resulting in poor visual contrast. Further, small-molecule imaging agents exclusively offer one detection mode, potentially limiting the use of a single contrast agent in the perioperative period. To overcome these pitfalls of the existing imaging agents, new imaging formulations with nanometer-scale dimensions are developed.^9^ These formulations permeate cancer tissues while sparing healthy tissues due to enhanced permeability and retention (EPR) effect or “passive targeting.”^10^ Additional surface functionalization of the nanoprobes (NPs) with tumor-targeting moieties such as antibodies, aptamers, nanobodies, and small molecules leads to an enhanced accumulation in cancer tissues, also known as “active targeting.”^11^ However, there exists an unmet need of integrating multiple optical imaging agents in a single nanoformulation that offers multiple complementary options for tracking throughout the perioperative time window. Additionally, upon the localization in tumor tissues, targeted stimuli-activated therapies (e.g., photothermal therapy, photodynamic therapy) can be performed using the same agent to enhance the versatility of these nanoformulations.^12^

DNA functionalized nanoparticles have shown enormous potential as cancer theranostic agents in the last decade.^13^ The inherent programmability of DNA enables designer nanomaterial development with numerous shapes, sizes, and functionalities. Recently, DNA-based nanomaterials have been used for biomedical applications such as cancer theranostics.^14-15^ Further, diverse custom DNA synthesis and conjugation chemistry allow the functionalization of DNA with other existing nanostructures, e.g., plasmonic/magnetic/silica/semiconductor nanoparticles, coordination polymers, lipid vesicles, liposomes, proteins, and organic fluorophores.^16^ Due to their inherent biocompatibility, DNA functionalized gold nanoparticles (AuNPs) are emerging as effective cancer imaging and therapeutic agents.^17-18^ AuNPs, specifically nanorods, possess inherent optoacoustic contrast due to the strong absorption of NIR light.^19^ Subsequent functionalization of DNA strands with fluorophores and MRI active moieties augment additional imaging modalities.^17, 20-21^ Recently, a dual-modality nanoparticle with SERS and fluorescence imaging capabilities was developed using DNA-based optimization strategy and applied for cancer imaging and therapy.^22^

Herein, we have developed a generalized fabrication strategy of AuNP-based triple modal nanoprobes (TMNPs) with three complementary optical modalities: optoacoustic, fluorescence, and surface-enhanced Raman Scattering (SERS) with complementary strengths. Optoacoustic modality is amenable for localization of deep-seated tumors before surgery but suffers from low sensitivity and cannot be used during surgery.^23^ On the other hand, fluorescence is detected using a fluorescence camera during the surgery, thus guide surgeons to excise the tumor.^24^ Yet, the use of this modality is limited because of photobleaching of fluorophores and autofluorescence from surrounding normal tissue.^25^ The SERS modality offers much-improved specificity than fluorescence due to “fingerprint”-like spectral features.^26-28^ However, the time-consuming spectral acquisition for a large tumor volume is not suitable for real-time image-guided surgery, although several rapid-Raman systems are under development.^29^ Recently, several examples of SERS NP-based cancer imaging have showcased near-perfect delineation of tumors in vivo due to the specificity of the Raman signatures from the NPs.^30-32^ We postulated that an amalgamation of optoacoustic, fluorescence and SERS modalities would provide a platform for preoperative localization of malignancies using optoacoustic, intraoperative guidance using fluorescence, and clean margin verification using SERS. However, a challenge exists in reconciling SERS and fluorescence, whose intensities are competing functions of the average nanoparticle-surface-to-fluorophore distance. While SERS intensity is highest near the surface, the fluorescence signal from the same molecule is quenched due to the proximity.^33-35^ While it is challenging to control fluorophore to Au surface distance, DNA molecules are utilized as a nanoscopic ruler to place fluorophores to Au surface in nanometer-scale precision.^36-37^ Herein, we leveraged the programmability of DNA functionalized nanoparticles for the fabrication of functional TMNP with three complementary optical imaging modalities, viz. optoacoustic, fluorescence, and SERS, for targeted imaging of breast cancer in vivo. We functionalized the TMNP outer surface to improve the specificity of targeting the FOLR1 positive TNBC. Further, we use these TMNPs for image-guided photothermal ablation of the tumor tissue.

## Results and Discussion

### Design Considerations for TMNPs

Current fabrication methodology allowed the fabrication of TMNP with any arbitrary shape Au nanoparticles, such as spherical AuNP (50 nm), Au nanocube (40 nm edge), Au nanostar (15 nm core, 10 nm spikes), and Au nanoprism (60 nm edge) (Figure S1). However, we preferred gold nanorod (AuNR) as the plasmonic component due to efficient interaction with NIR light, thanks to localized surface plasmon resonance (LSPR) maxima around 780 nm. Further, AuNRs have been shown to enhance SERS signal intensity in the NIR excitation window and demonstrate better photothermal therapy (PTT) efficacy.^38^ In a typical synthesis procedure, Hexadecyltrimethylammoniumbromide (CTAB) coated AuNRs are functionalized with 5’ thiol-ended DNA strands with different lengths with a red fluorophore attached to the 3’ end (See method section in the SI for details). After purification, positively charged NIR fluorophores were incubated in a 100:1 ratio to DNA-coated AuNRs for 4 hours. The nanoprobes were further coated with a silica layer to enhance the biostability (Figure 1a). The surface of the TMNPs was functionalized with folate groups to improve the targeted delivery to TNBC overexpressing FOLR1. The plasmonic AuNR cores produced an optoacoustic signal upon excitation by pulsed NIR laser (680-900 nm, 10 ns pulse duration) in a multi-spectral optoacoustic tomography (MSOT) imaging system. The NIR fluorophore emitted fluorescence emission with NIR (680-750 nm) excitation. Further, NIR fluorophores also produced SERS photons upon excitation with a NIR laser (785 nm) due to the proximity of the AuNR surface and were detected by a confocal Raman microscope. Plasmon-induced heating of TMNPs upon the exposure of NIR laser (660 nm) leads to selective ablation of tumor tissue (Figure 1b). The red fluorophores were used for in vitro TMNP uptake studies.

**Figure 1.**
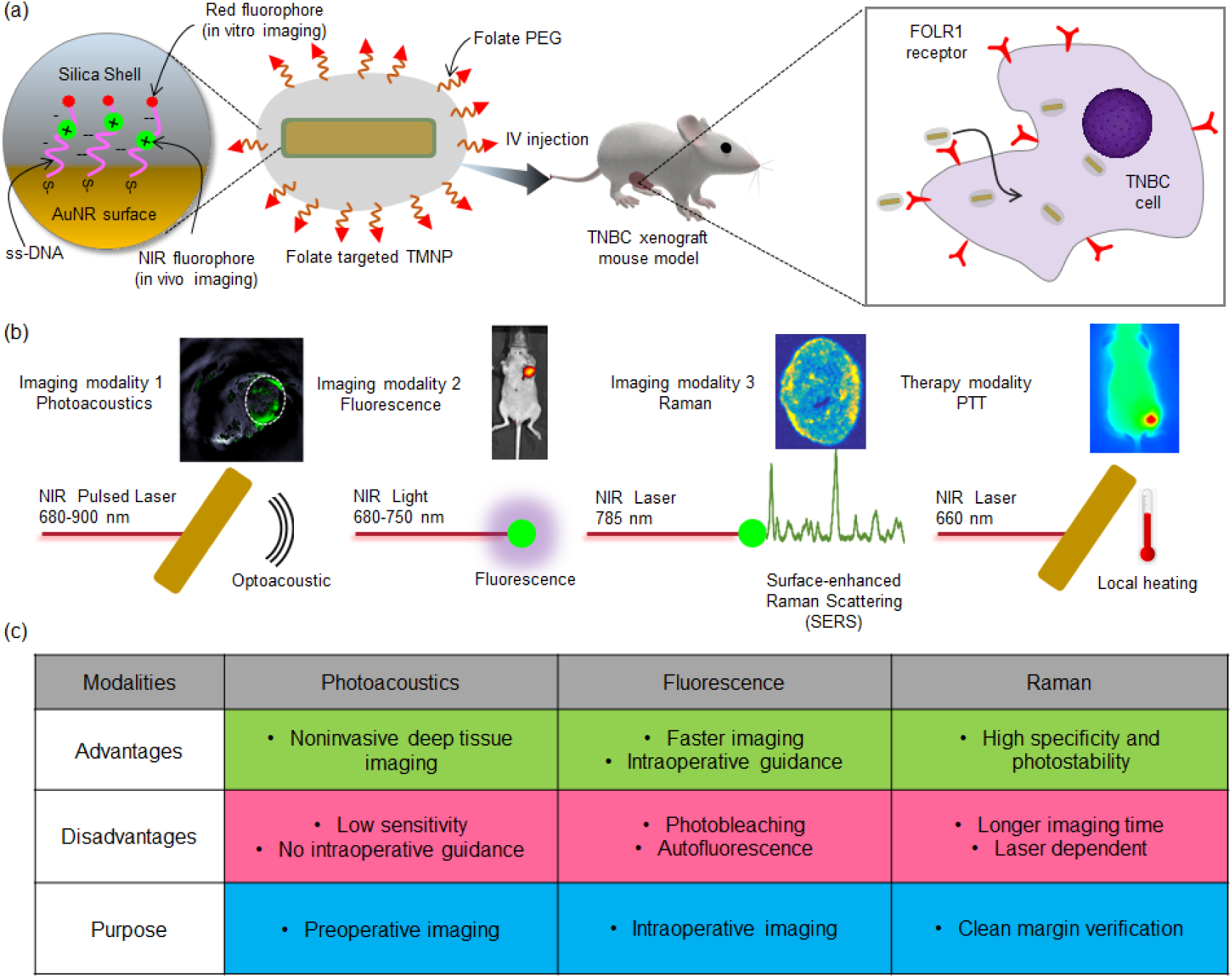
(a) Schematic representation of the TMNP. 5’ end of ss-DNA chains is attached to the AuNR surface via a thiol-Au bond. The positively charged NIR fluorophores are attached to negatively charged DNA via electrostatic interaction and a red fluorophore is attached to the 3’ end of the ss-DNA for in vitro fluorescence imaging and flow cytometry. A silica layer was formed around the DNA-capped AuNRs and further functionalized with PEG_5000_-folate groups. The folate-targeted TMNPs are used for in vivo multimodal targeted imaging of TNBCs overexpressing FOLR1. (b) Different optical imaging and therapy modalities offered by the TMNPs for in vivo imaging: (i) Optoacoustic signal is obtained when a NIR pulsed laser is shined on the AuNR, (ii) A detectable fluorescence light and SERS signal are emitted from the NIR fluorophore when excited by NIR light and laser, respectively. (iii) NIR light also induce plasmon mediated local heating leading to specific killing of cancer cells. (c) Advantages and drawbacks of three imaging modalities and the incorporation of three modalities ensures the monitoring perioperative time window.

### DNA-based Optimization of SERS Signal

The crucial parameter that influences the SERS enhancement factor is the distance of the SERS reporter from the nanoparticle surface. The average surface-fluorophore distance increases with increasing shell thickness, leading to a diminished SERS signal due to a lower electromagnetic field enhancement. We conducted extensive molecular dynamics (MD) simulations to prove this hypothesis. We chose poly-thymine sequences for minimal interaction with the Au surface, which permits maximum DNA loading.^39^ To simplify the simulation, we vertically tethered three single-stranded (ss) poly-thymine sequences (T_5_, T_10_, T_15_) on a gold surface via a thiol group, maintaining a surface coverage of one DNA/10 nm^2^. Four copies of Cy7 molecules were included in the simulation box. The MD simulation results suggested the fluorophores started to come in proximity to the gold surface in the presence of a DNA shell due to an electrostatic attraction force. The average distances of the fluorophores from the surface were proportional to the DNA shell thickness (Figure 2a). Based on this theoretical cue, we designed the four different DNA shells-T_5_, T_10_, T_15_, T_20_ (Figure 2b) and derived nanoprobes using IR 780 perchlorate NIR fluorophore (See SI for details). We found that the SERS intensity, after subtraction of the fluorescence background, was proportional to the DNA shell thickness. The intensity at 953 cm^-1^ was ∼18.9, 8.5, and 3.9 times higher for T_5_, T_10_, T_15_ shells, respectively, compared to the T_20_ DNA shell (Figure 2c, Table S1). Next, we measured the NIR fluorescence intensity at 800 nm from the TMNPs derived from T_5_, T_10_, T_15_, and T_20_ shells. On the other hand, the fluorescence intensity from T_20_, T_15_, and T_10_ coated TMNPs at 800 nm was ∼6.9, 3.8, and 1.3 times higher than that of T_5_ DNA functionalized TMNPs, respectively (Figure 2d).

**Figure 2.**
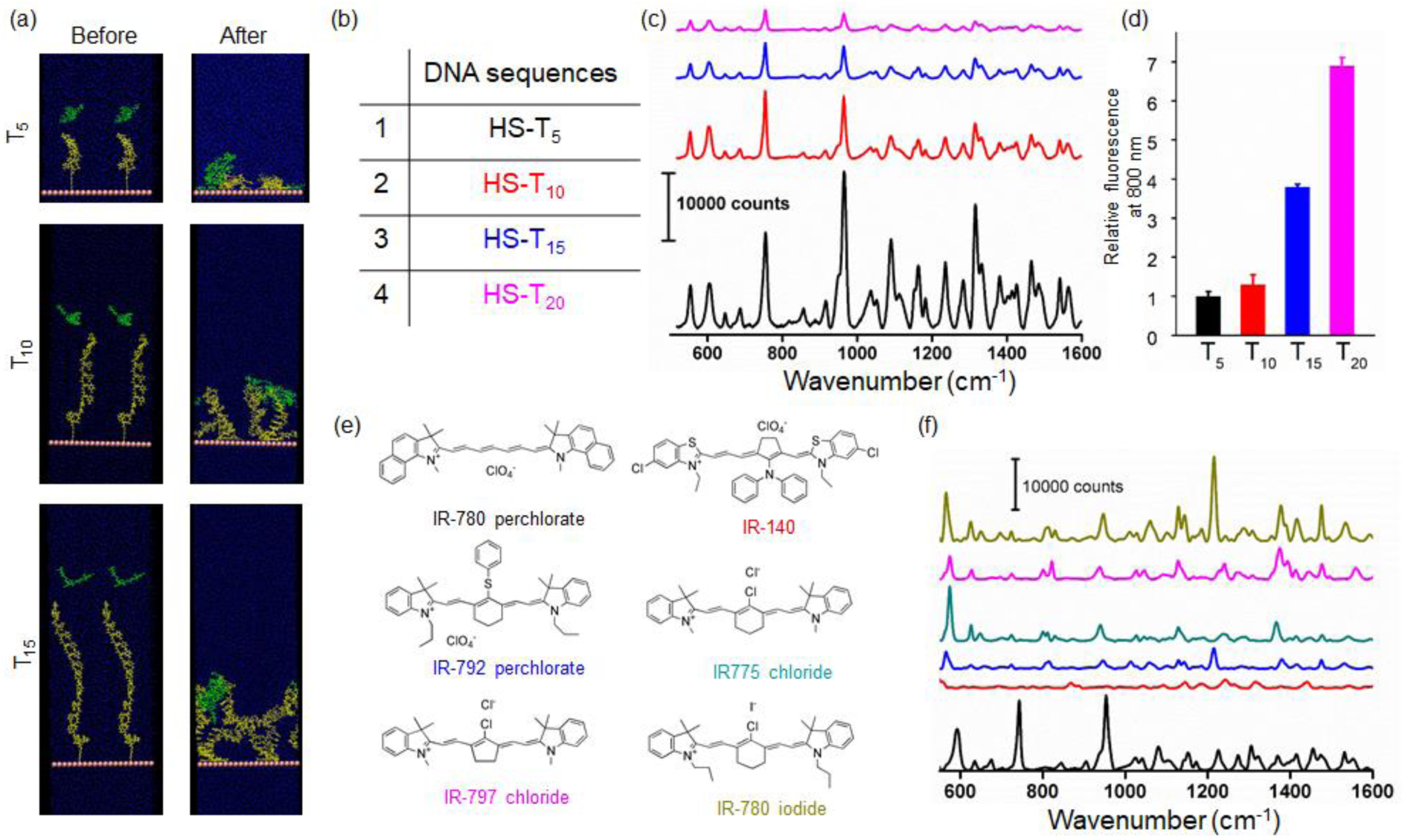
(a) Snapshots of T_5_, T_10_, T_15_ DNA sequences (yellow chain) and fluorophore (green) at the beginning and end of MD simulations in water. With increasing chain length, the average fluorophore-AuNR surface distance increase. (b) Four different sequences that are used for this study. (c) Fluorescence background-subtracted SERS spectra of 10 fM solutions of TMNP derived from T_5_ (black), T_10_ (red), T_15_(blue), T_20_ (purple) DNA showing a higher SERS spectrum for T_5_ DNA-capped TMNPs. (d) Relative fluorescence intensity of 10 fM solutions of TMNP derived from T_5_ (black), T_10_ (red), T_15_(blue), T_20_ (purple) show significant quenching of the fluorescence from the TMNPs. (e) Chemical structures of six NIR fluorophores that are used for this study. (f) Fluorescence background-subtracted SERS spectra of 10 fM solutions of TMNP derived from T_10_ DNA with six NIR fluorophores showing unique spectra for all the fluorophores. Screening of fluorophores determines that IR-780 perchlorate and IR-780 perchlorate is the brightest.

Next, we aimed to improve the SERS signal output by screening different commercially available NIR fluorophores. It is well established that different fluorophores possess different Raman scattering cross-sections at the same excitation wavelength. We selected six commercially available, positively charged NIR fluorophores with different molecular structures (Figure 2e). All the NIR fluorophores possessed a positive charge that allowed the surface attachment towards negatively charged DNA-coated AuNRs. We used the T_10_ DNA coated AuNR under the same concentration of SERS reporters, and fabricated the six TMNPs and measured the intensities of six TMNPs at 10 fM concentrations. All the TMNPs show their unique Raman signatures, and IR-780 perchlorate and IR780 iodide were exhibited the brightest Raman signal (Figure 2f, Table S2).

### Characterizations of TMNPs

The limit of detection for the IR780 perchlorate was determined to be ∼5 fM, which is comparable to several existing SERS nanoprobes for in vivo cancer imaging (Figure S2). Based on the effect of DNA-shell on the Raman and fluorescence signal intensities, TMNPs with T_10_ DNA with IR-780 perchlorate fluorophores provided the optimum fluorescence and Raman signal and was selected for in vivo imaging purposes. After synthesizing TMNPs, we prepared a plastic pouch containing 200 µL of 10 fM TMNPs and imaged it in MSOT, fluorescence, and Raman imaging systems emulating the in vivo imaging conditions. All three imaging platforms independently imaged the nanoprobes (Figure S3). We observed less than 10% decrease in the SERS and fluorescence signals after 24 hours serum incubation under physiological conditions, possibly due to degradation of the silica shells. These results suggested excellent in vitro stability of fluorescence and SERS signals. Furthermore, TMNPs did not exhibit any significant cytotoxicity measured by WST1 assay towards breast cancer cells after 24 hours post-incubation (Figure S4).

### In vitro and In vivo FOLR1 Targeting Using TMNPs

These encouraging results prompted us to perform targeted multimodal imaging of TNBC using the TMNPs as the contrast agent. Recently, FOLR1 has emerged as a promising biomarker for TNBCs.^40^ The silica surface of the TMNPs was further modified to append folate-PEG_5000_ groups using well-known NHS-ester chemistry to synthesize folate-targeted nanoprobes (Figure 3a) to enhance the specificity and tumor uptake. We measured the hydrodynamic diameter to be 96±38 nm with a particle tracking analyzer system. Transmission electron microscopy (TEM) confirmed that the majority of TMNPs were a single AuNR surrounded by the silica shell (Figures 3b, S5). To demonstrate the FOLR1 targeting capability, we selected two TNBC cell lines, namely, MDA-MB-231 and MDA-MB-468. We recently demonstrated that MDA-MB-231 and MDA-MB-468 display a low and high FOLR1 expression, respectively, and folate targeted DNA origami nanostructures were preferentially endocytosed in MDA-MB-468 cells.^41^ First, we incubated both the cell lines with 10 fM folate targeted TMNPs at 37 °C for 4 hours. We observed a higher uptake of the TMNPs in the MDA-MB-468 cells compared to MDA-MB-231 cells, indicating a FOLR1 mediated uptake of the targeted nanoparticles (Figure 3c). We used flow cytometry-based measurements to quantify the targeting, which exhibited ∼7.2 times higher uptake in the MDA-MB-468 cells than MDA-MB-231 cells (Figure S6). The successful in vitro targeting results encouraged us to develop a bilateral TNBC xenograft mouse model by inoculating MDA-MB-468 cells and MDA-MB-231 cells on the left and right back side of nude mice, respectively. We intravenously administered TMNPs (200 µL, 10 nM) in the bilateral human BC xenograft mouse model (n=3). 24 hours post-injection, we imaged mice under a small animal imaging system under 750 nm NIR excitation light. We observed that the fluorescence intensity from the left tumor was higher than that of the right tumor (Figure 3d). We further excised both the tumors, imaged ex vivo, and quantified the fluorescence intensity. The left tumor was ∼2 times brighter than the right tumor (Figure 3e). We carried out immunohistochemistry for both the tumors, which revealed that FOLR1 was overexpressed in the left tumor compared to the right tumor (Figure 3f). These results suggested that TMNPs were efficiently targeting FOLR1 positive TNBC in vitro and in vivo. Nonetheless, in vivo contrast was found to be less than in vitro targeting as FOLR1 low expressing MDA-MB-231 tumors could uptake TMNPs via the passive targeting or EPR effect.

**Figure 3.**
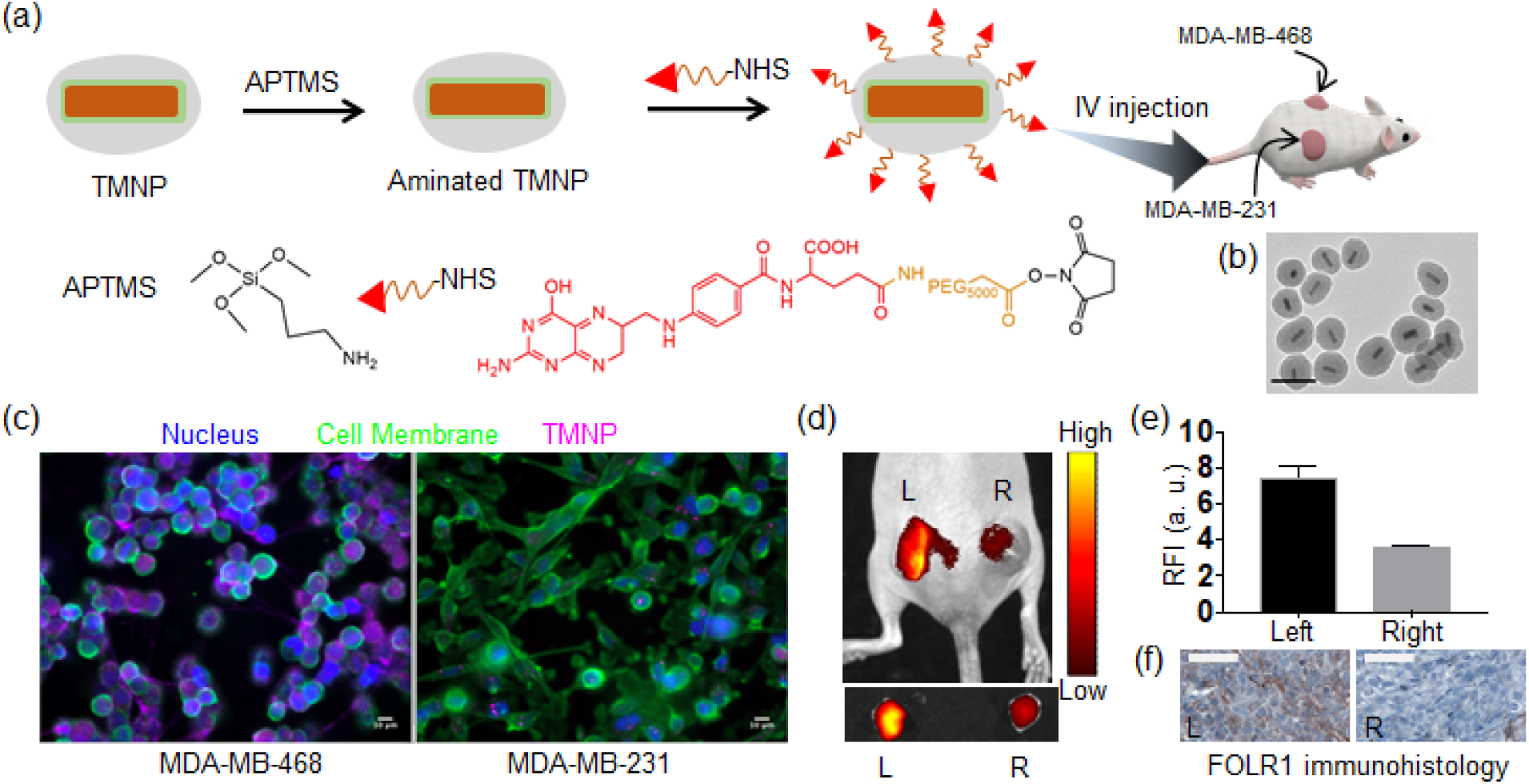
(a) Functionalization of TMNP with Folate-PEG_5000_ moieties: First, TMNPs were aminated with (3-Aminopropyl)trimethoxysilane (APTMS) and thereafter reacted with folate-PEG_5000_-NHS ester. The folate-targeted TMNPs were used for assessing in vitro and in vivo targeting. (b) Transmission electron microscope (TEM) image of the folate functionalized TMNPs. Scale bar is 100 nm. (c) In vitro FOLR1 targeting: widefield epifluorescence microscopy imaging of MDA-MB-468 cells and MDA-MB-231 cells incubated with 10 fM TMNPs show an enhanced uptake in FOLR1 overexpressing MDA-MB-468. Cell nucleus and membrane were stained with Hoechst^®^ 33342 and wheat germ agglutinin, AF488 conjugate respectively. Scale bars are 10 μm. (d, e) NIR fluorescence image of TNBC xenograft mouse 24 hours after being intravenously injected with TMNPs (200 µL,10 nM). In this mouse, MDA-MB-468 and MDA-MB-231 cells are inoculated on the left and right sides, respectively. The in vivo and ex vivo imaging of tumors reveal a significantly higher fluorescence. (f) FOLR1 immunohistology of the left and right tumors show FOLR1 overexpression in the left tumor. Scale bars are 50 μm.

### Multimodal Imaging of TNBC-patient Derived Xenograft Mouse Model Using TMNPs

It has been confirmed that FOLR1 is expressed inhomogeneously in TNBC tissue due to the inherent heterogeneity in protein expression.^40, 42-44^ We further verified the overexpression of FOLR1 in TNBC tissues compared to normal breast tissue in a breast cancer tissue array. FOLR1 immunohistochemistry of TNBC tissues confirmed an inhomogeneous FOLR1 expression throughout the tissue (Figure S7). To demonstrate the multimodal imaging and FOLR1 targeting capabilities of TMNPs, we used TNBC patient-derived xenograft mice (n=3) that were intravenously injected TMNPs (200 µL, 10 nM). We imaged the live mice in a multispectral optoacoustic tomography (MSOT) device before and 24 hours post-injection. Using multispectral unmixing, the signal from TMNPs could be differentiated from major intrinsic absorbers, namely, hemoglobin (Hb) and Oxyhemoglobin (HbO_2_) in 680-800 nm window (Figure S8). 24 hours post-injection, we observed a significant increase in the TMNP-specific MSOT signal from the tumor area (Figure 4a). Next, we imaged the same mice using NIR excitation of 750 nm and observed intense fluorescence light from the tumor area (Figure 4b). The fluorescence-positive area correlates with the bioluminescence emanated by the luciferase tagged TNBC patient-derived xenograft in the live mice (Figure 4c). We excised tumor tissue and tumor-adjacent normal tissue using fluorescence light guidance and imaged it under a Raman microscope. Tumor tissue exhibited the Raman signature of TMNPs, while normal tissue did not contain any such signature, verifying our observations in the fluorescence imaging (Figure 4d). Next, we interrogated if the TMNPs could capture the heterogeneous overexpression of FOLR1 in the tumor tissue. Extensive H&E histology and FOLR1 immunohistology of the excised tissue suggested FOLR1 expression heterogeneity throughout the tumor (Figures 4e, f). For example, in the periphery (green box), FOLR1 was overexpressed compared to the internal region (for example, orange box). We carried out point-by-point high-resolution spectral acquisition using a confocal Raman microscope. The fluorescence intensity map at 825 nm revealed an inhomogeneous fluorescence intensity throughout the tissue (Figure 4g). After the fluorescence background subtraction, a Raman intensity map at 953 cm^-1^ was created (Figure 4h), showing different Raman intensity levels throughout the tissue. The average fluorescence intensity at 825 nm and Raman intensity at 953 cm^-1^ of the FOLR1 overexpressing region (green box) was ∼4.7 times higher than that of FOLR1 lower expressing region (orange box) (Figures 4i, j). These investigations strongly suggested that folate-targeted TMNPs accumulated in FOLR1 overexpressing regions in TNBC. Atomic absorption spectroscopy (AAS)-based biodistribution studies of the post mortem mice also revealed accumulation of TMNPs in the tumor, along with other RES organs like the liver and spleen, 24 hours post-injection (Figure S9).

**Figure 4.**
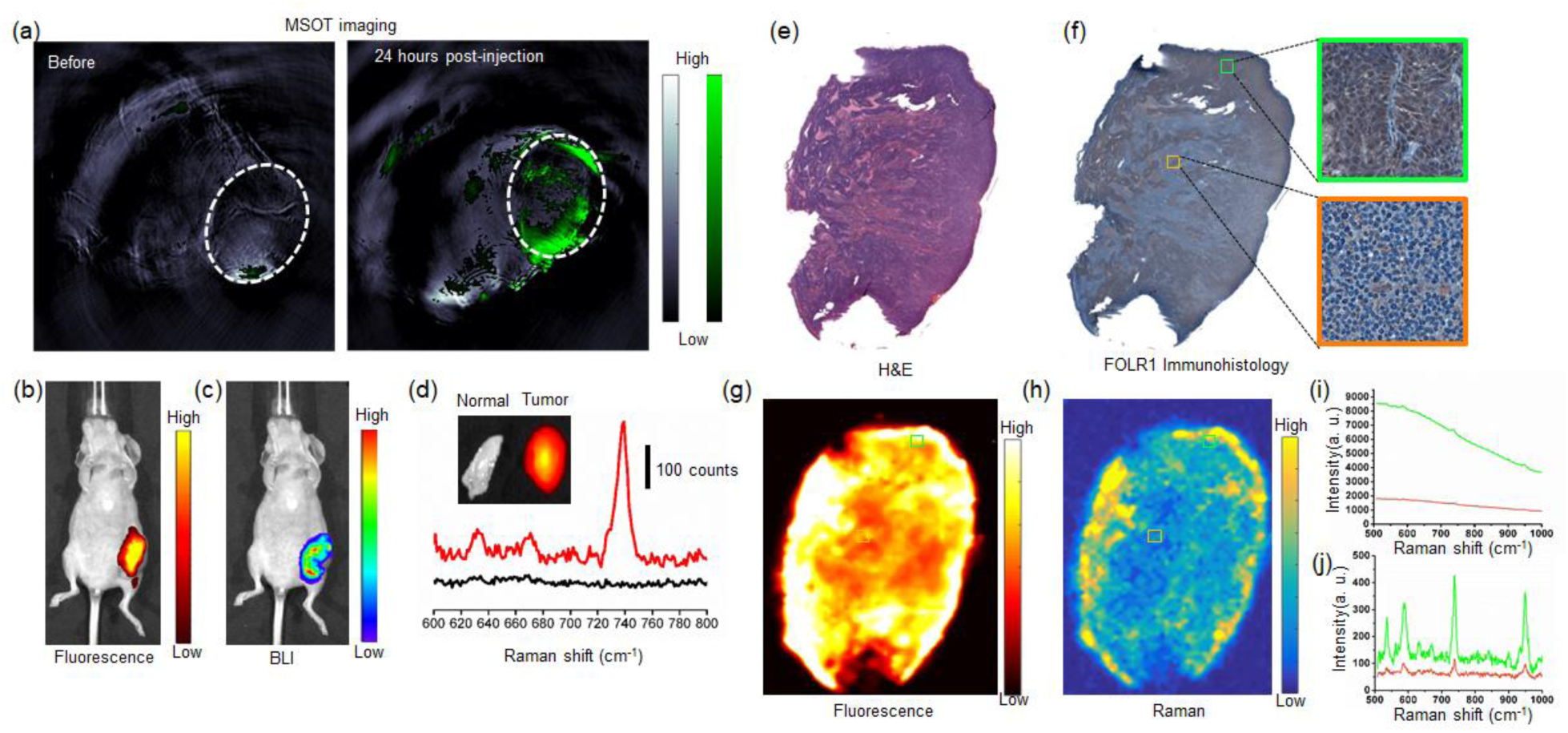
(a) MSOT images of the PDX mouse before and 24 hours after TMNP injection show an improved contrast in the tumor area. (b, c) NIR fluorescence image of the same mouse showing a significantly higher fluorescence from tumor area, which corresponds very well with the bioluminescence (BLI) images of the mouse. (d) Ex vivo fluorescence images and background subtracted Raman spectra of the tumor (red) and tumor-adjacent normal tissue (black), showing a significantly higher fluorescence and SERS signal from the tissue. (e, f) H&E and FOLR1 immunohistology of the excised tissue show intratumor heterogeneity of the FOLR1 expression. (g, h) The high-resolution fluorescence image and Raman image of the same tissue show heterogeneity of fluorescence and SERS intensity. Average spectra of the green box and orange box acquired in a confocal Raman microscope: (i) without fluorescence background subtraction (ii) with fluorescence background subtraction showing a higher fluorescence and Raman intensity in the tissue overexpressing FOLR1.

### In vivo PTT of Using TMNPs

Encouraged by the selective accumulation of TMNPs in the tumor tissue in vivo, we performed image-guided PTT. We established an orthotopic TNBC xenograft mouse model with MDA-MB-468 cells on both sides of the back of nude mice (Figure 5a). We injected TMNPs (200 µL, 20 nM) for the treatment group and 200 µL 10 mM PBS for the control group, both 6 days after inoculation (n=5/group). We waited for 24 hours to allow the TMNPs to accumulate in the tumor. After verifying the presence of TMNPs with NIR fluorescence imaging, we irradiated the right tumor of both the groups with a 660 nm laser (power density of 3 W/cm^2^) for 5 minutes. We also measured the temperature increase with a thermal camera which showed the temperature in the TMNP injected mouse was ∼20 °C higher than that of PBS injected mouse (Figures 5b, c). We monitored the tumor growth every 4 days after the PTT. The average size of the right tumor of the TMNP-injected mice was significantly smaller than that of the left tumor of the same mouse. We did not observe any difference in the tumor size between the PBS injected mice left and right tumor, and TMNP injected mice the left tumor (Figures 5d, e). We did not observe any change in the body weights in the PBS and TMNP injected groups (Figure S10). Additionally, there were no histological differences in the major organs in these two groups suggesting no systemic toxicity was presented by the injection of TMNPs (Figures 5f, S10).

**Figure 5.**
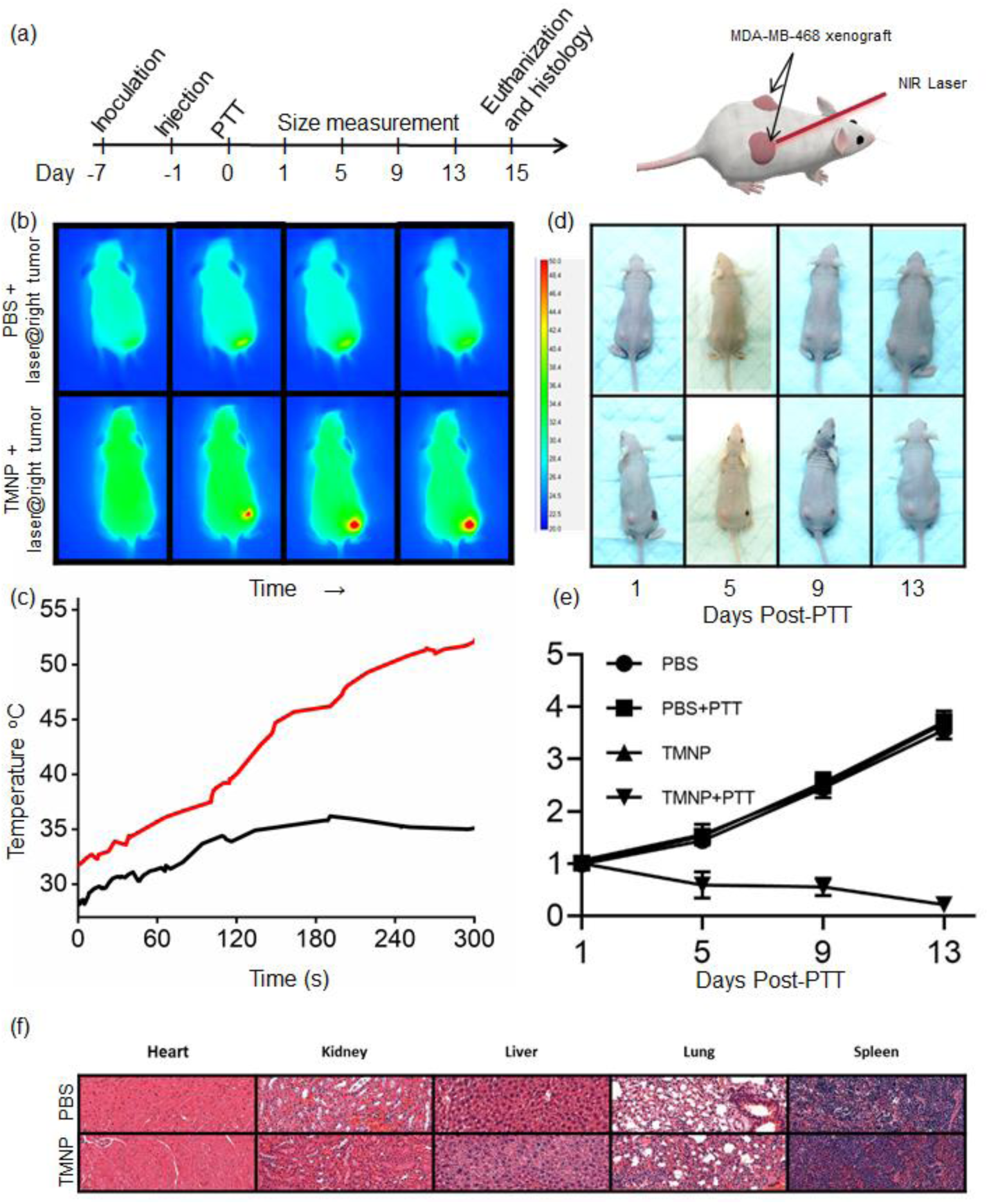
(a) The timeline for PTT. Mice were inoculated with MDA-MB-468 cells on both sides of the back. After six days, mice were injected with TMNPs, and the day after was irradiated with a NIR laser on the right tumor. The tumor size was measured on days 1, 5, 9, and 13, and on day 15, mice were sacrificed, and histology was performed in major organs. (b) Temperature maps of mice during PTT using a temperature camera for PBS injected (top), and TMNP injected mouse (below). (c) The average temperature of the tumor area for PBS (black) and TMNP (red) injected mice shows an increase in temperature for TMNP injected mice. (d) Representative photos of PBS injected (top) and TMNP injected mouse (below) post PTT show the TMNP injected mice’s right tumor decreased in size and disappeared after two weeks. (e) The change in the tumor size for all different groups proves the PTT induced destruction of tumor growth for TMNP injected mice.

## Conclusions

We have developed a fabrication strategy for versatile TMNPs that incorporates optoacoustic, fluorescence, and SERS as imaging modes. The presence of AuNR in the structure makes it amenable for PTT after particle localization in the tumor. The TMNPs offer deep-tissue imaging capability due to optoacoustic modality, real-time intraoperative visualization of cancer tissue using fluorescence light, and clean margin verification with high precision using SERS. DNA functionalization on the AuNRs offered unprecedented design flexibility in fluorophore selection and optimizing fluorescence and Raman signal intensity from the TMNPs. By changing the length of DNA coating on the AuNR, we optimized the fluorescence and SERS signals from the same fluorophore. We functionalized the surface of the TMNPs with folate molecules to target FOLR1 overexpressed in TNBC. We show successful targeting of TMNPs to the FOLR1 in TNBC cell xenograft mouse model. We also demonstrated the multimodal imaging capability of the TMNPs in TNBC patient-derived xenograft mouse models. Overall, the TMNPs accumulated preferentially in the FOLR1 overexpressing tissue. Using selective exposure of the tumor tissue with NIR laser, we found that the growth of TNBC was reversed due to an excellent therapeutic efficacy. We strongly believe these nanoprobes could be efficiently used for multiplexed real-time imaging of cancer biomarkers in vivo. The DNA-based strategy developed in this study would be particularly important for creating multimodal nanoprobes with renally excretable Au nanoclusters for achieving better chances of clinical translation.^45-47^

## Experimental Section

### Materials

All chemicals of the highest purity were purchased from Sigma-Aldrich (St. Louis, MO) and used without further purifications. All DNA sequences were purchased from IDT DNA Inc. as HPLC grade. G-25 size-exclusion columns were purchased from GE Healthcare. Folic acid-PEG-NHS of molecular weight of 5000 Da were purchased from Nanocs Inc (Cat. No. PG2-FANS-5k). Anti-Folate Binding Protein/FBP antibody (ab67422) was purchased from Abcam for immunohistochemistry. Human breast cancer tissue array purchased from US Biomax, Inc (catalogue numbers BC00432, BR1202).

### Gold Nanorod Synthesis

The synthesis of AuNR was done following the silver-assisted growth procedure described elsewhere.^48^ First gold seeds were synthesized by adding 60 µL of 10 mM ice cold NaBH_4_ solution to 1 mL of 2.5 mM HAuCl_4_ solution in 100 mM Hexadecyltrimethylammoniumbromide (CTAB) and vortexed vigorously. The solution color immediately turned to yellowish brown color. To synthesize the gold nanorods, to 100 mL of 10 mM HAuCl_4_ solution in 100 mM CTAB, 250 µL of 1 mM AgNO_3_ solution was added. After mixing, 70 µL of 80 mM ascorbic acid solution was added and mixed thoroughly. To this mixture 12 µL of the previously prepared AuNP seed solution was added and kept undisturbed for several hours. The solution turned purple indicating formation of AuNRs. The dimensions were measured using TEM imaging.

### Fluorophore Labelling of DNA

All the DNA were purchased as 5’ thiol and 3’ amine modifications. Fluorophore labeling of DNA was carried out by reacting CF™ 680R succinimidyl ester with 3’ amine-modified DNA. 100 μL of 10 mM NHS modified fluorophore solution in anhydrous N,N-dimethylformamide (DMF) was mixed with 100 μL of 1 mM amine modified DNA in DI water. To this reaction mixture, 20 μL of trimethylamine was added. After overnight reaction at room temperature, the reaction mixture was lyophilized and 100 μL, 10 mM Tris(2-carboxyethyl)phosphine hydrochloride (TCEP) solution was added to reduce S-S bond. The excess fluorophore and TCEP molecules were removed using a Zeba Spin desalting column. The final concentrations were measured using a micro-UV-Vis spectrophotometer using the extinction coefficient at 260 nm.

### Fabrication of Folate Targeted TMNPs

The concentrations of AuNPs were measured using nanoparticle tracking analysis (NanoSight NS300, Malvern Panalytical). As synthesized Au nanorod was centrifuged (13,000 rpm for 15 min) and washed with DI water 2 times to remove excess CTAB. The AuNRs were dispersed in 0.5% SDS and the fluorophore modified DNAs were added in 1:1000 ratio. A 5 M NaCl solution was slowly added in seven aliquots to bring the final Na^+^ concentration to 350 mM over 24 h. Then the solution was allowed to incubate overnight at room temperature. The excess DNA was removed by 3 times centrifugation (13,000 rpm for 15 min) and re-suspension in 1× TBE buffer. To a 100 μL 50 nM DNA functionalized AuNRs, 1 μL 0.5 mM NIR fluorophore solution was added and kept for incubation for 4 hours. To form the silica shell 13 mL isopropanol, 500 μL tetraethylorthosilicate (TEOS), and 200 μL of ammonium hydroxide (NH_4_OH) were added to 1.3 mL of 10 nM AuNR solution, in a 50 mL Falcon tube. The mixture was mixed in a shaker for 12 min. The mixture was centrifuged (13,000 rpm for 10 min) and washed with ethanol for 3 times.

The silica coated AuNRs were further functionalized with folate-PEG in a series of steps. In the first step, the silica surface was functionalized with amine. To silicated NPs (1 mL, 10 nM) in ethanol, 100 μL of 3-aminopropyltrimethoxysilane (APTMS) was mixed at room temperature for 1 hour. After that, the particles were washed 2 times with ethanol and 1 time with DI water. The aminated NPs were functionalized with folate using a Folic acid-PEG-NHS ester. To 1mL, 10 nM TMNP in 10 mM sodium bicarbonate buffer (pH 8.6), a 100 μL of 10 mM Folic acid-PEG-NHS solution was added and kept for shaking for 2 h in the dark at room temperature. Next, the particles were washed with water 3 times and redispersed in 10 mM PBS buffer (pH 7.4) for in vitro and in vivo studies.

### Molecular Dynamic (MD) Simulation

Computer simulation was performed to explore the fluorophore-DNA interactions with gold surface in aqueous media. We simulated 5-mer, 10-mer and 15-mer of single-stranded poly-thymine (T) sequences (thereby labeled as T_5_, T_10_ and T_15_) at their all-atom representation. In each cases, the initial configuration of the simulated setup consisted of 4 copies of DNA sequences vertically tethered with a gold surface via sulfur group linker (-S-C_6_-), maintaining one DNA/ 10 nm^2^ surface coverage. Four copies of Lumiprope Cy7 molecules were included in the simulation box. The simulation box was explicitly solvated with water and counterions were added to keep the solution charge-neutral. AMBER-99bsc11 force fields were used for modeling DNA.^49^ AMBER-DYES2 forcefield for fluorophore and TIP3P3 model to represent water molecules were used.^50-51^ To investigate the role of DNA molecules in fluorophore-surface interaction, we also performed additional control simulations of surface adsorption dynamics of fluorophore in absence of DNA molecules. In these cases, the simulation setup was very similar to the original system, except that the DNA chains were absent.

Equilibrium molecular dynamics simulation was employed for sampling the adsorption process using GROMACS-20184 software package (www.gromacs.org). Each of the systems were subjected to 100 ns MD simulation and two independent realizations of each of the simulations were carried out by varying the initial velocity distributions. An integration time step of 2 femtosecond was used. All simulations were performed in NVT ensemble. The simulation temperature was maintained at 300.15K via V-rescale thermostat.^52^ LINCS algorithm was used to constrain all the hydrogen bonds involving DNA and fluorophores.^53^ The water molecules were kept rigid using SETTLE algorithm.^54^ Verlet cutoff scheme was employed throughout the simulations with Lennard Jones interaction extending to 1.2 nm.^55^ Particle mesh Ewald summation was used for treating the electrostatic interactions.^56-57^ The gold surface was kept restrained in all three directions such that no force is being calculated. This is achieved by using ‘freezegrps’ option within GROMACS.

### Cell Culture

All cell lines were purchased from American Type Culture Collection (ATCC; Manassas, VA, USA). MDA-MB-231 (ATCC HTB-26) and MDA-MB-468 (ATCC HTB-132) cell lines were maintained in folate-free Dulbecco’s Modified Eagle’s Medium (DMEM) media supplemented with 10% foetal bovine serum (FBS), 100 U mL^−1^ penicillin and 100 µg mL^−1^ streptomycin at 37 °C in 95% humidified air with 5% CO_2_.

### Cell Viability Studies

WST1 assay: WST-1 assay was carried out using WST1 (Roche) reagent using the specification of the manufacturer. Briefly, ∼10,000 MDA-MB-468 cells were seeded in a sterile 96-well microplate. The cells were incubated with TMNPs (500, 50, 5 fM) for 24 hours. For positive control, cells were treated with 70% Methanol for 2 h. After the incubation, 10 μL of WST1 reagent was added. After 2 h of incubation at 37 °C, the UV–Vis absorbance of the well was measured at 440 nm. The intensity at 440 nm was compared with the untreated control to determine the cell viability.

### In vitro Uptake of TMNPs

Flow cytometry-based TMNP uptake studies were carried out by seeding 1 × 10^5^ MDA-MB-468 and MDA-MB-231 cells in 6-well plates. After 24 hours, growth medium was replaced with a new medium containing 10 fM targeted TMNPs. After 4 hours, cells were detached with 0.25% trypsin and 0.05% EDTA in 10 mM PBS. The cells were washed twice with 10 mM PBS before it was measured using a flow cytometer (BD LSR II flow cytometer, BD Biosciences) using the CF™ 680R fluorescence signal. Dead cells are excluded using propidium iodide (PI) staining. All flow cytometry data were analyzed by FlowJo software. For fluorescence imaging, after 4 hours cells were fixed in 4% paraformaldehyde in 10 mM PBS and stained with Hoechst® 33342 (1 μg/ml, for nucleus) and wheat germ agglutinin AF488/633 (5 μg/ml, for cell membrane) before imaging in widefield fluorescence microscope (Zeiss, Axioplan 2).

### Animal Studies

Human xenograft mouse models were carried out under the protocol number 06-07-011 approved by the Institutional Animal Care and Use Committees (IACUC) at Memorial Sloan Kettering Cancer Center. We have complied with all relevant ethical regulations.For subcutaneous breast cancer tumors, 8-week-old female outbred homozygous nude mice (Foxn1nu, Jackson Laboratory) were subcutaneously injected with 2 × 10^6^ MDA-MB-231 (ATCC HTB-26) and MDA-MB-468 (ATCC HTB-132) cells suspended in 0.2 mL culture media mixed with 0.2 mL of Matrigel (Corning) into the right and left lower back sides, respectively. We waited for 2 weeks post inoculation for in vivo imaging experiments. For PTT, both the sides were inoculated with MDA-MB-468 (ATCC HTB-132) cells and waited for 6 days before the TMNP injection.

The PDX of triple-negative breast tumor was established using triple-negative breast cancer bone metastasis specimens. Freshly resected tumor was also diced into 1–2 mm pieces in 10 cm sterile culture dishes in serum-free MEM medium supplemented with non-essential amino acids and antibiotics. The tissue pieces were transferred to 24-well plate and transduced with infected cell supernatants of lentiviral vectors expressing GFP-luciferase described elsewhere.^46^ Tumor growth was periodically monitored by bioluminescence imaging by IVIS system by retro-orbitally injecting 100 mg kg^−1^ d-luciferin (14681, Cayman Chemical).

### Multimodal In vivo Imaging

MSOT imaging: We used pre-clinical multi-spectral optoacoustic tomography (MSOT) device (MSOT inVision 256, iThera Medical, Munich, Germany). The equipment was furnished with an array of 256 detector elements which are cylindrically focused, having a central ultrasound frequency of 5 MHz and up to 270° coverage. MSOT imaging was performed in a temperature-controlled water bath at 34 °C. Spatial reconstruction of the data was performed using the ViewMSOT software suite (V3.6; iThera Medical) and a back-projection algorithm. Fluorescence imaging: Mice were administered 200 μL of 10 nM TMNPs in 10 mM PBS buffer via tail vein injection. Mice were anesthetized with isoflurane for in vivo imaging or euthanized with CO_2_ and then dissected for ex vivo imaging. Fluorescence images were acquired with an IVIS-200 imaging system (Xenogen Corp., Hopkinton, MA). Radiance (photons/s^−1^ cm^−2^) was calculated for the region of interest using in built Living Image V4.2 software.

Raman imaging: All Raman scans were carried out using an InVia Raman microscope (Renishaw). The microscope was equipped with a piezo-controlled stage, a 300 mW 785 nm diode laser and a 1-inch CCD detector with a spectral resolution of 1.07 cm^−1^. The SERS images were acquired through a ×5 objective lens (Leica). Typically, Raman scans were carried out at 100 mW laser power, with 1.5 s acquisition time, using the StreamLine high-speed acquisition mode. All Raman images were acquired and analyzed under the same conditions, including the same laser power, Raman integration times, and focal volume (same objective lens). For the analysis of the acquired images, a graphical user interface using MATLAB (R2014b) and PLS Toolbox v.8.0 (Eigenvector Research, Inc., Wenatchee, WA, USA) developed in-house was used.

### Biodistribution of TMNPs

Homogenized tissue specimens of known mass from euthanized mice injected with TMNPs (n = 3) were put in Savillex PFA vials. 1 mL of freshly prepared aqua regia (1:3 conc. nitric acid and hydrochloric acid) to the vials and heated for 12 h at 60 °C for complete digestion. When the samples fully dissolved in the acidic solution, 100 μL of the digest were taken up in 15 mL Falcon tubes and the volume was made up to 5 mL using deionized water. The samples were analyzed with an atomic absorption spectroscopy (Atomic Absorption Spectrometer Analyst 800) calibrated with known HAuCl_4_ concentrations.

### Histology

After imaging, multiorgan specimens were fixed in 4% paraformaldehyde (MP Chemicals, Solon, OH, USA) overnight at 4 °C, followed by a rinse with PBS for 15 min, and then kept in 70% ethanol until embedding in paraffin. Paraffin embedding was performed using ASP6025 automatic tissue processor (Leica Biosystems) and sectioned 5 μm thickness using a Leica RM2265 automated paraffin microtome (Leica Biosystems). Sections were stained with haematoxylin and eosin (H&E) for histopathological analysis and scanned with Mirax digital slide scanner (Zeiss). Images were analyzed with Pannoramic Viewer software (3DHistech).

## ASSOCIATED CONTENT

### Supporting Information

The Supporting Information is available free of charge. Supporting figures S1-10, supporting tables 1,2.

## AUTHOR INFORMATION

### Author Contributions

S.P and M. F. K. conceived and designed the project. The experimental work has been performed by S.P. with the help from C.A., M.W., T.R. V. K. R. provided PDX mouse model. C.A. developed the data analysis interface. S.P. and C.A. performed data analysis. J.K.K. and J.M. carried out MD simulations. All authors contributed to discussions on the project. The manuscript was written through contributions of all authors. All authors have given approval to the final version of the manuscript.

### Notes

Dr. Moritz Kircher, unfortunately passed away during the preparation of the manuscript. The authors declare no competing financial interest.

## ACKNOWLEDGMENT

The authors thank the MSKCC electron microscopy and molecular cytology core facilities for technical support. We acknowledge NIH R01 EB017748, Startup Research Grant from SERB (SRG/2019/000953), Innovative Young Biotechnologist award by DBT (BT/12/IYBA/2019/14) and Research Initiation Grant from IIT Bhilai. TR thanks SERB (CRG/2019/007013) for funding. JM acknowledges support of the Department of Atomic Energy, Government of India, under Project Identification No. RTI 4007. We thank IIT Bhilai for providing research infrastructure.

