## Supplemental Information for "DNA-functionalized Gold Nanorods for Targeted Triple-modal Optical Imaging and Photothermal Therapy of Triple-negative Breast Cancer"

| DNA Shell | Absolute Intensity at 953 cm^-1^(cps) | Relative Intensity (a. u.) |
| --- | --- | --- |
| T_20_ | 2580 | 1.0 |
| T_15_ | 10108 | 3.9 |
| T_10_ | 22035 | 8.5 |
| T_5_ | 48924 | 18.9 |

Supplementary Table 1

| NIR fluorophore  (CAS number) | Maximum Raman Peak Position (cm^-1^) | Absolute Intensity (cps) | Relative Intensity (a. u.) |
| --- | --- | --- | --- |
| IR 780 Perchloate (23178-67-8) | 953 | 7602 | 8.3 |
| IR 140 (53655-17-7) | 1242 | 916 | 1.0 |
| IR 792 Perchlorate (207399-10-8) | 1214 | 2115 | 2.3 |
| IR 775 Chloride (199444-11-6) | 574 | 5540 | 6.0 |
| IR 797 Chloride (110992-55-7) | 1375 | 3159 | 3.4 |
| IR 780 Iodide (207399-07-3) | 1215 | 8509 | 9.3 |

Supplementary Table 2

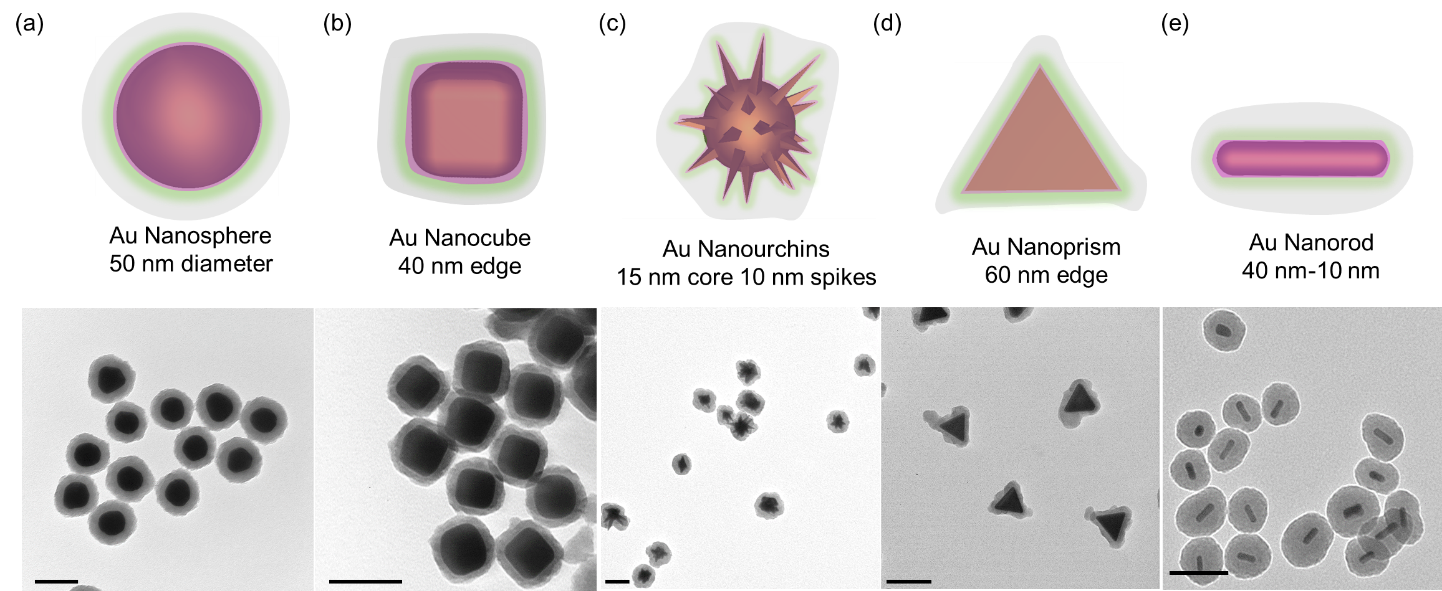

**Figure S1:** TEM images of TMNPs synthesized using (a) 50 nm Au nanosphere, (b) Au nanocube with 40 nm edge, (c) Au nanourchins with 15 nm core and 10 nm spikes, (d) Au nanoprism with 60 nm edge, (e) Au nanorod with 40 nm length and 10 nm width. Scale bars represent 100 nm.

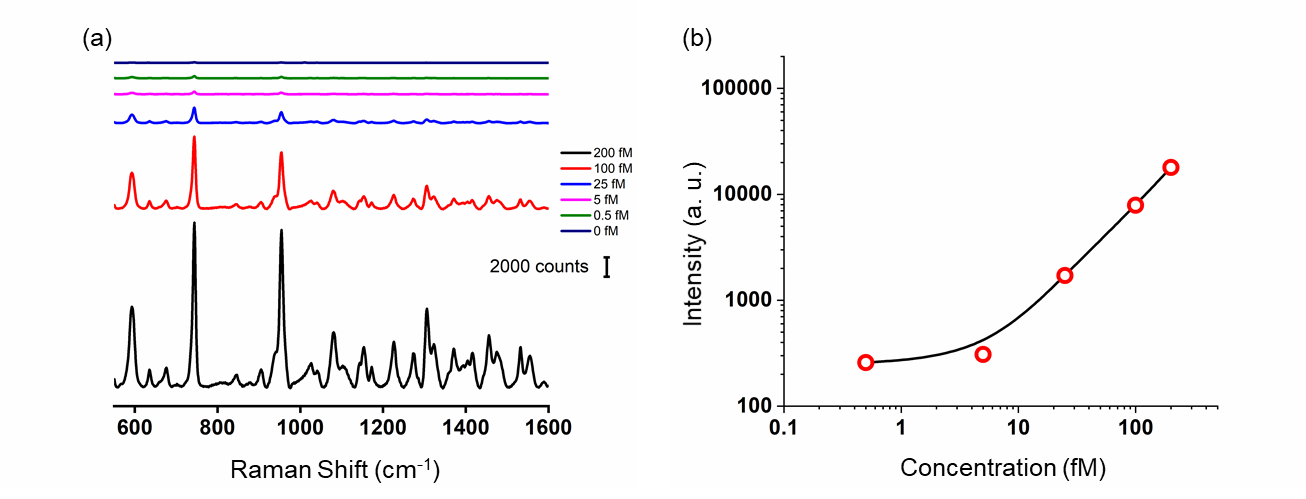

**Figure S2:** Limit of detection of the optimized TMNPs. (a) Background subtracted SERS spectra of TMNPs with decreasing concentration in 50% serum using the same imaging conditions used for in vivo imaging. (b) The limit of 5 fM concentration TMNPs was determined to be ~5 fM.

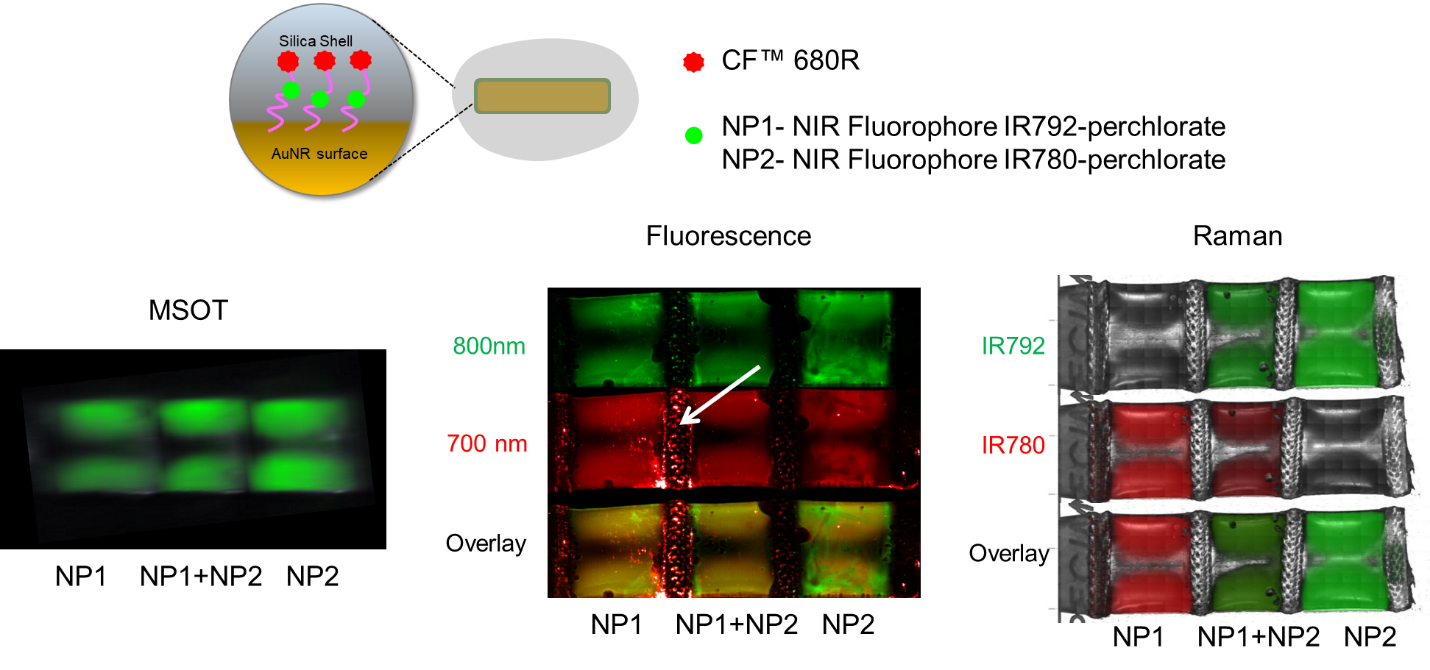

**Figure S3:** Multimodal imaging of the 200 µL of 10 fM TMNPs diluted in 50% serum in plastic pouch. NP1 and NP2 are fabricated using IR 792 and IR 780, respectively. Both the TMNPs carried CF 690R. Using all the imaging methods, TMNPs were detected. Strong nonspecific fluorescence in 700 nm channel was found (white arrow) due to the autofluorescence.

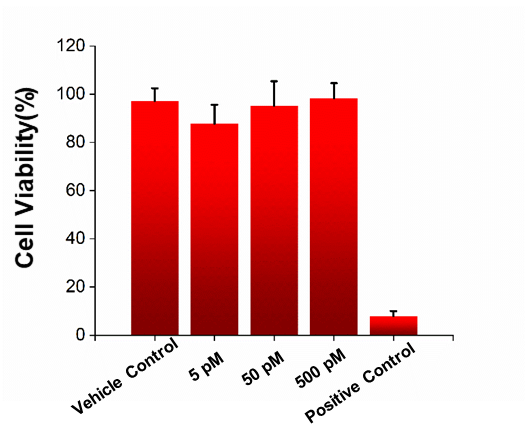

**Figure S4:** WST1 cell viability assay of MDA-MB-468 cells incubated with 500, 100, and 50 pM of OFRNPs showing no cell toxicity of the TMNPs.

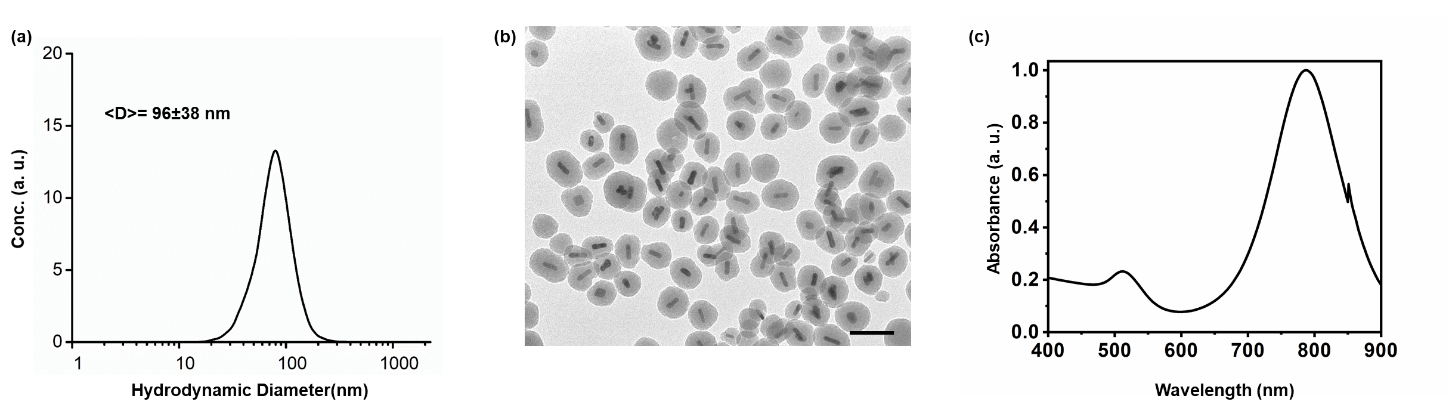

**Figure S5:** (a) Hydrodynamic diameter distribution of the folate-PEG coated targeted TMNPs obtained from nanoparticle tracking measurement. (b) TEM images of TMNPs. (c) UV-Vis spectra of the TMNPs showing strong absorbance around 780 nm.

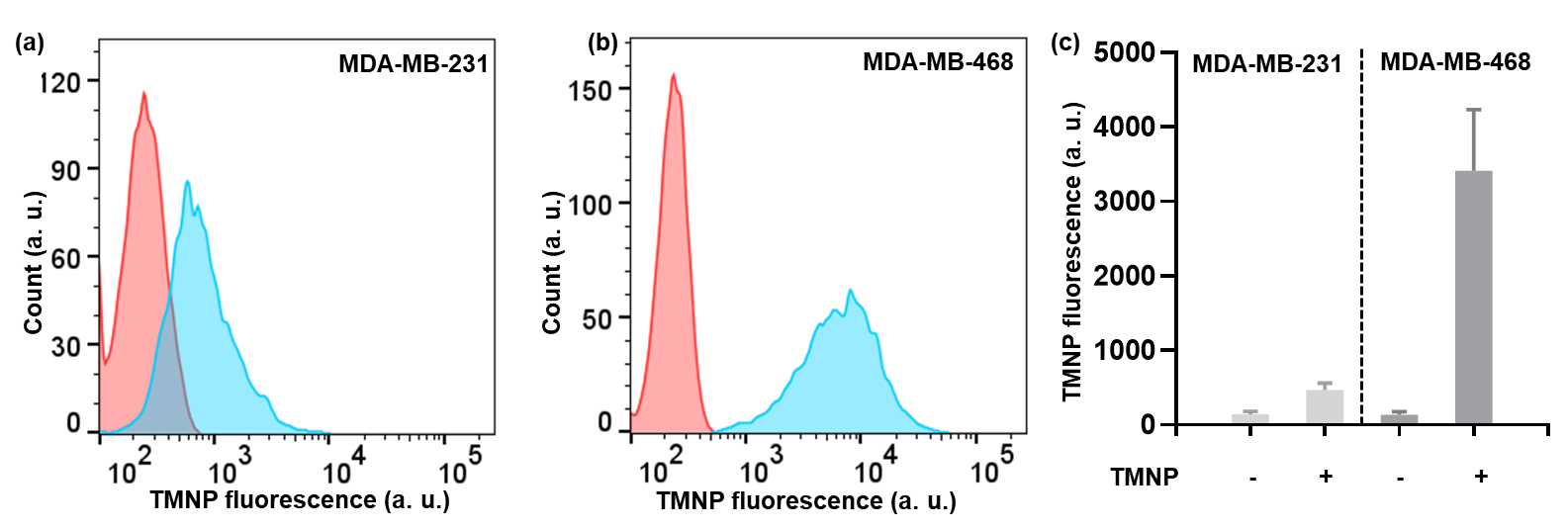

**Figure S6:** Flow cytometry-based uptake assay of TMNPs in TNBC cells. (a, b) MDA-MB-231 cells and FOLR1 overexpressing MDA-MB-468 cells incubated with 10 fM TMNPs (cyan) or without any treatment (red) at 37 °C for 4 hours. (c) Quantification of fluorescence intensity show a significant uptake of folate targeted TMNPs in MDA-MB-468 cells. Fluorescence from CF 680R fluorophore was used for this assay.

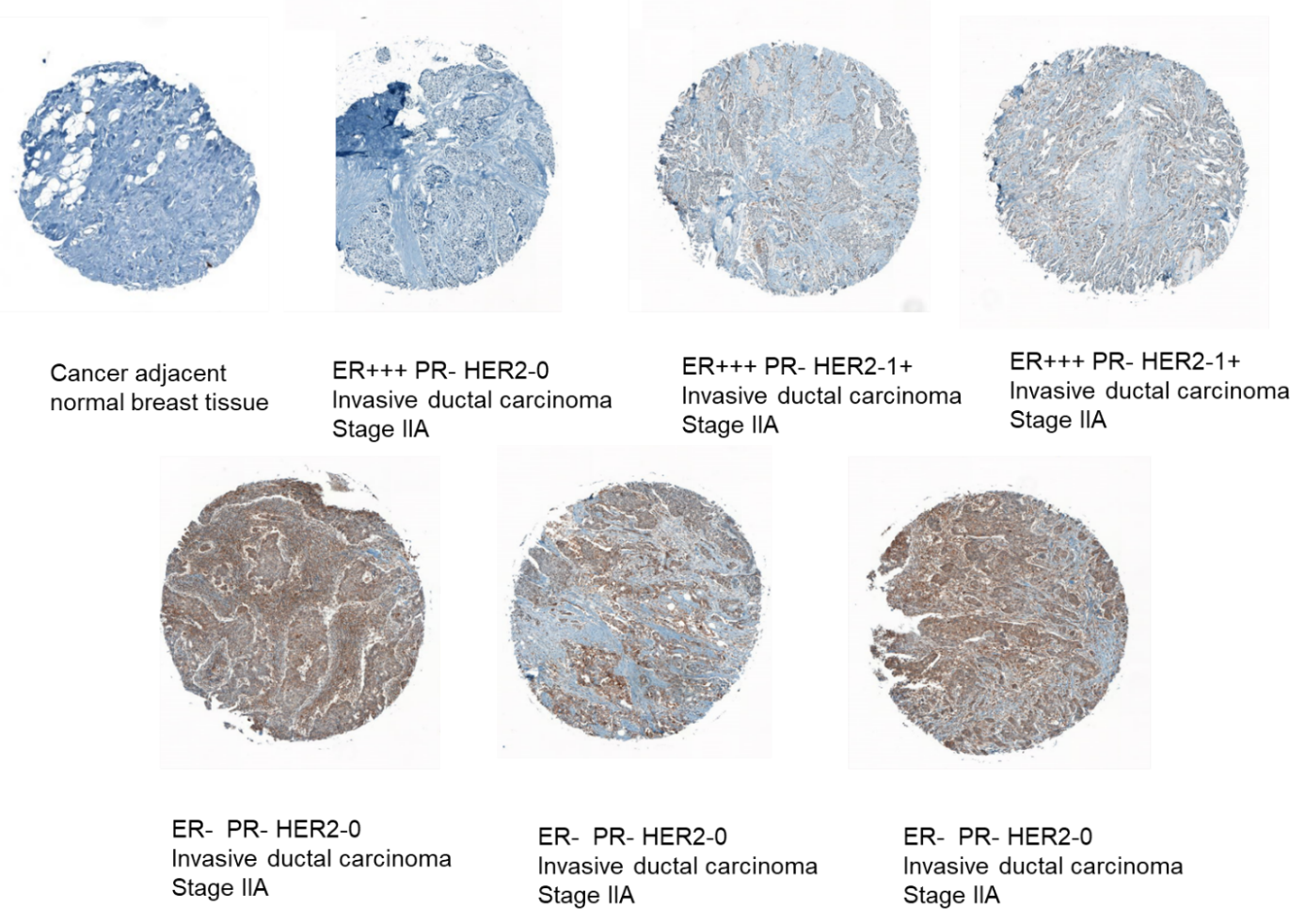

**Figure S7:** FOLR1 immunohistology of breast cancer of normal tissue in breast cancer tissue array. FOLR1 expression is relatively high in TNBC (bottom row) compared to other subtypes and cancer adjacent normal tissue. There is also intratumoral heterogeneity of FOLR1 expression.

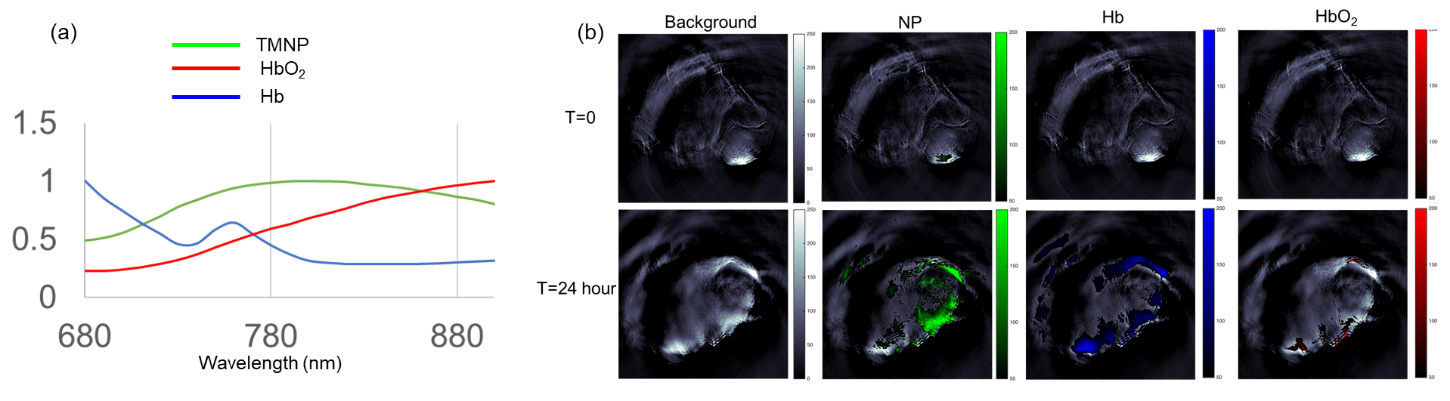

**Figure S8:** (a) Spectral features of TMNP (green), oxygenated hemoglobin (red), hemoglobin (blue) used for the deconvolution of the MSOT signal. (b) Corresponding map of TMNPs TMNP (green), oxygenated hemoglobin (red), hemoglobin (blue) shows an enhanced signal coming from tumor region due to the presence of TMNPs.

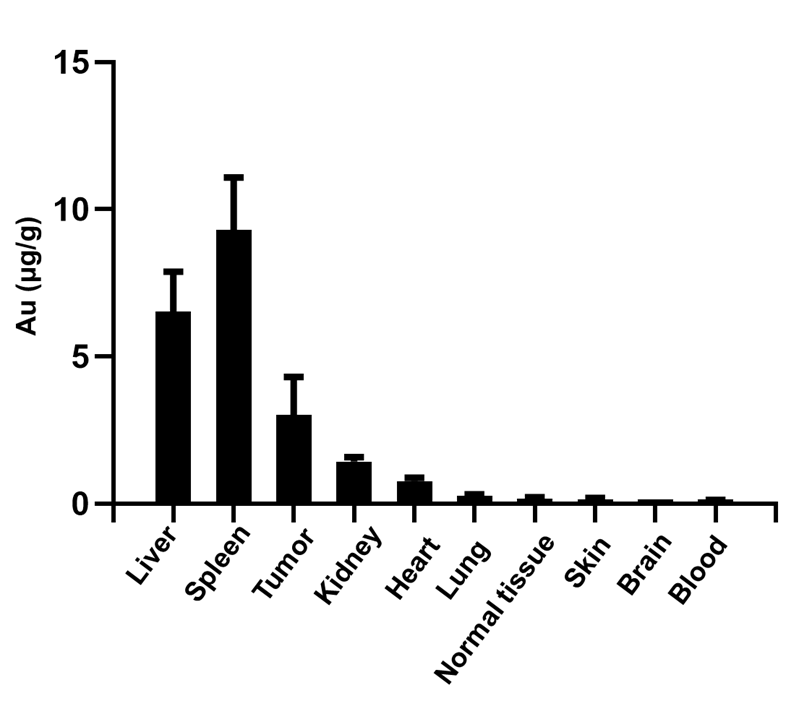

**Figure S9:** Biodistribution of TMNPs in TNBC PDX mice (n=3), 24 hours post-injection. measured by atomic absorption spectroscopy (AAS). RES organs like liver, spleen have high uptake. Tumor tissue demonstrates significantly higher localization of the NPs compared to normal tissue.

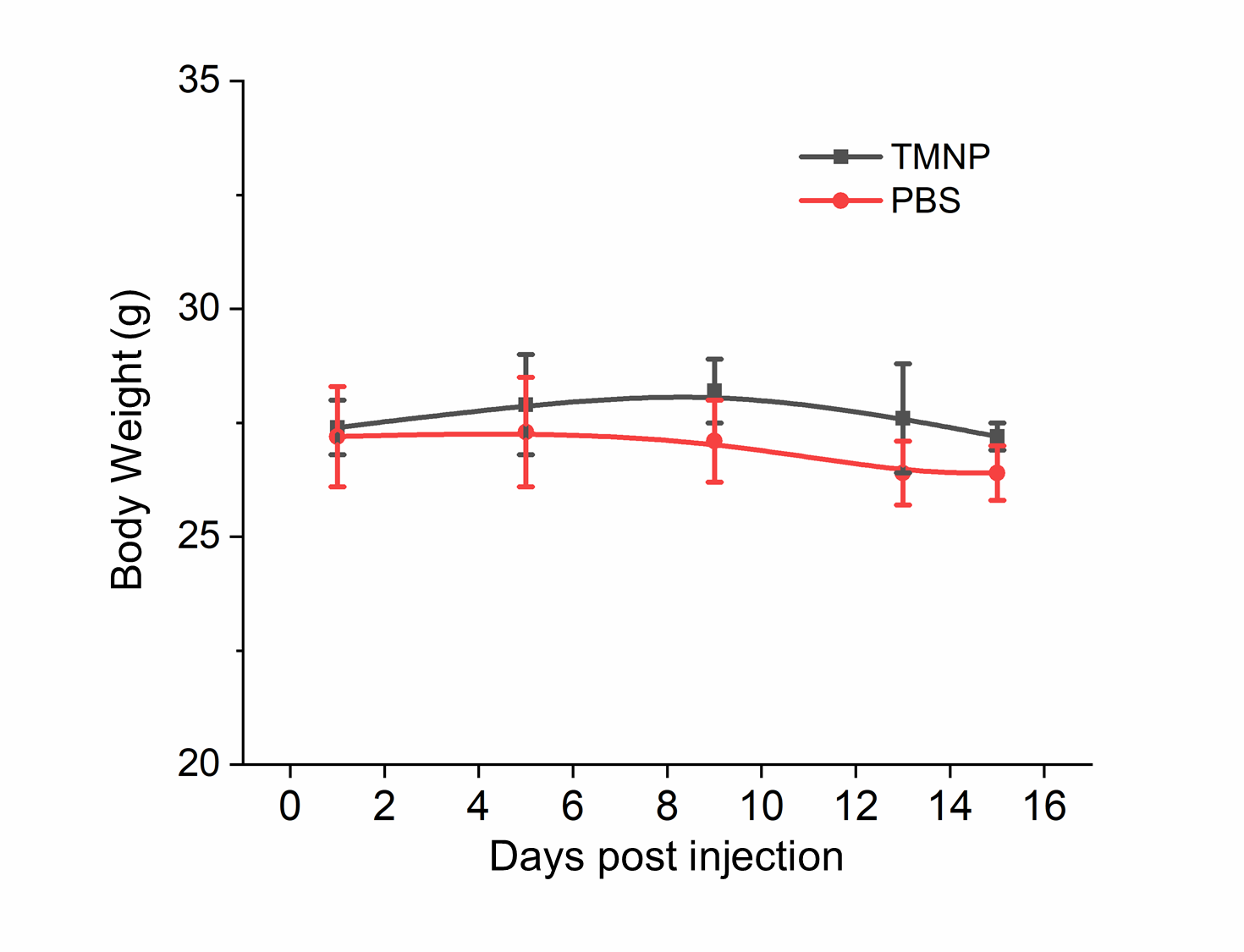

**Figure S10:** Body weight of TMNP (black) and PBS (red) injected groups (n=3) show no changes in body weight over 15 days.
